# A Pre-transplant Blood-based Lipid Signature for Prediction of Antibody-mediated Rejection in Kidney Transplant Patients

**DOI:** 10.1101/460030

**Authors:** Monther A Alsultan, Gaurav Gupta, Daniel Contaifer, Sindhura Bobba, Dayanjan S. Wijesinghe

## Abstract

There is a lack of biomarkers for pre-kidney transplant immune risk stratification to avoid over- or under-immunosuppression, despite substantial advances in kidney transplant management. Since the circulating lipidome is integrally involved in various inflammatory process and pathophysiology of several immune response, we hypothesized that the lipidome may provide biomarkers that are helpful in the prediction of kidney rejection. Serial plasma samples collected over 1-year post-kidney transplant from a prospective, observational cohort of 45 adult Kidney Transplant [antibody-mediated rejection (AMR)=16; stable controls (SC) =29] patients, were assayed for 210 unique lipid metabolites by quantitative mass spectrometry. A stepwise regularized linear discriminant analysis (RLDA) was used to generate models of predictors of rejection and multivariate statistics was used to identify metabolic group differences. The RLDA models include lipids as well as of calculated panel reactive antibody (cPRA) and presence of significant donor-specific antibody (DSA) at the time of transplant. Analysis of lipids on day of transplant (T1) samples revealed a 7-lipid classifier (lysophosphatidylethanolamine and phosphatidylcholine species) which discriminated between AMR and SC with a misclassification rate of 8.9% [AUC = 0.95 (95% CI = 0.84-0.98), R^2^ = 0.63]. A clinical model using cPRA and DSA was inferior and produced a misclassification rate of 15.6% [AUC = 0.82 (95% CI = 0.69-0.93), R^2^ = 0.41]. A stepwise combined model using 4 lipid classifiers and DSA improved the AUC further to 0.98 (95% CI = 0.89-1.0, R^2^ = 0.83) with a misclassification of only 2.2%. Specific classes of lipids were lower in AMR compared with SC. Serial analysis of SC patients demonstrated metabolic changes between T1 and 6 months (T2) post-transplant, but not between 6 and 12 (T3) months post-transplant. There were no overtime changes in AMR patients. Analysis of SC T1 vs AMR T3 (that at time of AMR) showed sustained decreased levels of lipids in AMR at the time of rejection. These findings suggest that lack of anti-inflammatory polyunsaturated phospholipids differentiate SC from AMR pre-transplant and at the time of rejection, and a composite model using a 4-lipid classifier along with DSA could be used for prediction of antibody-mediated rejection before transplant.

**Highlights:**

1. Despite significant advancements in kidney transplant treatment and intensive clinical follow-up monitoring, all rejection events are unlikely to be recognized at the beginning. As a result, efforts have been made to identify new biomarkers for kidney rejection detection.
2. While lipids are known to be potent mediators of inflammation, pro-resolving processes, and other cell signaling cascades, lipidomics can be applied to identify reliable biomarkers to monitor disease severity and may also allow prediction of kidney rejection.
3. Our lipidomic study shows lipid profile changes between antibody-mediated rejection group and stable control group as a function of different time point, pre and post-kidney transplantation. Furthermore, our study demonstrates that combining lipid and clinical parameters allow prediction of rejection on the day of the transplant.
4. These findings have the potential to change the present paradigm of pre and post-transplant monitoring and management of these patients by implementing an evidence-based risk stratification technique, resulting in a substantial improvement in kidney transplant success.

## 1-INTRODUCTION

End-stage renal disease (ESRD) affects almost 786,000 people in the United States in 2020, with 71% requiring dialysis and 29% requiring a kidney transplant^1^. Renal transplantation is a common therapeutic option for individuals with ESRD, and compared to dialysis, and it provides considerable improvements to quality of life and longer life benefits^2, 3^. Kidney transplant has advanced significantly over the last 50 years, beginning with the first successful organ transplant between twins in 1954^4^. Additionally, patient and allograft survival rates have surpassed 95% in the modern age of kidney transplantation^2^. In addition, patients who receive a kidney transplant have a much lower adjusted death rate (48.9 per 1,000) than those who receive dialysis (160.8 per 1,000)^1^.

Despite significant advancements in understanding the pathophysiology and immunology of most types of allograft rejection and kidney transplant treatment regimens, transplant rejection remains a significant concern and one of the leading indicators of long-term allograft loss^2, 5^. In the post-transplant monitoring for renal rejection, routine clinical follow-up evaluations are combined with laboratory testing of blood creatinine levels^6^. A combination of clinical (e.g., GFR, proteinuria), immunological (e.g., DSA), instrumental (e.g., resistive index at Doppler ultrasonography), and histological measures are being used to monitor the transplanted kidney. Overall, these “conventional biomarkers” have a number of limitations that are connected to disease as well as nephrologists’ and pathologists’ abilities^7^. A renal allograft biopsy for histological assessment is usually initiated when blood creatinine levels rise over a patient-specific baseline value, whether or not clinical symptoms arise. This frequently suggests detection in later stages of dysfunction, but early stages of dysfunction, when functional impairment is not yet clinically visible, may go unnoticed. As a result, protocol biopsies were developed to identify acute rejection in subclinical conditions^8^. Serum creatinine levels, glomerular filtration rate, and proteinuria are all monitored as part of the transplant monitoring process. These indicators are nonspecific; therefore, diagnosis necessitates an invasive allograft biopsy. Furthermore, because these markers have limited sensitivity for injury processes, this technique only detects pathological processes at a rather advanced stage of tissue injury and misses subclinical alterations. In recipients of kidney allografts, protocol biopsies have been proposed to identify alterations before graft failure is evident. Multiple biopsies, however, are required to diagnose subclinical alterations^6^. Although the invasive allograft biopsy, which is the gold standard for diagnosing and distinguishing between different forms of rejection and pathologic processes used to diagnose kidney rejection, has improved over time, hemorrhage and graft loss still occur after the procedure. Moreover, the practicality and cost of repeated biopsies required to capture anti-allograft immunity are major limitations, as sampling mistakes and inter-observer variability in biopsy findings^5, 9^. Even with repeated biopsies, it’s doubtful that all rejection episodes would be diagnosed at the outset, much alone the dangers and consequences that come with such a pricey procedure^8^. Furthermore, despite intense immunosuppressive medications and rigorous pre-transplant screening using donor-specific antibodies and degree of sensitization, kidney rejection is still poorly predicted, and 10-year survival rates have remained stable in recent decades^3, 10^. Immunosuppressive medication in kidney transplant patients is now administered according to protocols and adjusted based on the allograft’s functional or histological assessment, as well as indicators of drug toxicity or infection. As a result, a considerable percentage of patients are likely to get too much or too little immunosuppression, putting them at risk for infection, cancer, and medication toxicity, as well as acute and chronic graft harm from rejection^11^. Even measuring immunosuppressive medication levels in the blood and strict adherence to the specified medication levels does not preclude either overimmunosuppression, which can lead to infections, or underimmunosuppression which can lead to graft rejection or persistent immunological damage^10^. These drawbacks emphasize the need for more reliable, noninvasive approaches to detect and diagnose acute and chronic graft lesions earlier and more precisely^6^. As a result, one of the field’s main goals is to develop noninvasive indicators of allograft renal rejection^12^. Noninvasive biomarkers for transplant diseases have been developed as a result of these limitations. High-throughput molecular approaches have made it easier to find novel biomarkers that can help physicians regulate immunosuppression and predict problems and transplant success^13–15^.

Metabolomic seeks to assess all pertinent small molecules, enabling relative and absolute measurement of hundreds of lipids and water-soluble small molecules from low volumes of biological materials like blood. Observations of changed lipids concentrations reveal functionally meaningful read-outs of disturbed disease-associated pathways in human metabolism as intermediate and endpoint indicators of numerous biologic processes in the human body^12^. This type of global profiling promises to be very beneficial for discovering new prognostic and diagnostic indicators^16^. Endogenous lipids and their metabolites, which are important regulators of inflammation, pro-resolving activities, and other cell signaling cascades, are quantified via lipidomic profiling. Lipidomics has been used to demonstrate lipid homeostasis abnormalities in a variety of disorders, including immunological responses that produce inflammation^17^, such as kidney rejection. Importantly, these lipid alterations appear to occur at precise points throughout illness development, implying that lipidomics can be used to discover meaningful biomarkers for disease severity monitoring. In this regard, the ability to read the lipidome could be a useful tool for the risk stratification and prediction of kidney rejection after transplantation^17, 18^. Thus, the objectives of this study are to identify lipid biomarkers that could allow for better risk stratification to enhance the benefit and limit the risk of kidney rejection as well as enhance the immunosuppression therapy strategies for individuals. In addition, our objectives are to better understand the molecular pathophysiology underlying the development and progression of kidney rejection using lipidomic analysis.

## METHODS

### Patient Selection

This study was approved by the Virginia Commonwealth University Institutional Review Board (IRB). Patients were selected from a prospective observational cohort of a single-center adult kidney transplant center in the United States. The study population consisted of 16 consecutive patients who developed antibody-mediated rejection within 2 years of kidney transplant and 29 stable control (SC) patients who did not develop rejection at any point of post-transplant follow-up. Serial plasma samples are collected and stored at Time 1 (T1 - pre-transplant), Month 6 (T2) and Month 12(T3) and then yearly for all patient’s post-transplant as part of an IRB approved biobank protocol at our institution.

SC patients were selected based upon convenience sampling. The primary determinants for inclusion of SC patients was availability of volume of samples for lipid-based research assays and long-term follow-up with stable renal function with absence of rejection and known adherence to immunosuppressive regimens. A minimum follow-up of 2 years was mandated to be a candidate for inclusion in the study. Pediatric kidney recipients and multi-organ transplant recipients were excluded from the study.

### Immunosuppression and Treatment of Antibody-mediated Rejection

All patients received induction with anti-thymocyte globulin (Thymoglobulin, Genzyme, Cambridge, MA) with a total of 6 mg/kg over four consecutive days beginning in the operating room. Maintenance immunosuppression included a combination of tacrolimus, mycophenolate mofetil and prednisone tapered to 5 mg/day. Highly sensitized patients received 6 sessions of pre-emptive plasmapheresis with intravenous immunoglobulin (IVIG; 100mg/kg) based upon a pre-specified protocol as reported by us previously^19^.

Indication biopsies were performed for acute allograft dysfunction defined as a rise in creatinine >20% above baseline, serum creatinine nadir ≥2.0 mg/dL post-transplant; or delayed graft function >21 days post-transplant. Surveillance biopsies were performed in patients with a positive flow-cytometric crossmatch (T or B >100 mean channel shifts) and/or presence of pre-formed donor-specific antibody [DSA; >5000 mean fluorescent intensity (MFI)] at 1 month and 6-month post-transplant. Biopsies were graded based upon the Banff criteria^20^. Patients with AMR were treated with 6-9 sessions of plasmapheresis with intravenous immunoglobulin (IVIG; 100mg/kg) in conjunction with intravenous methylprednisolone 500mg administered once daily for 3 days. In selected cases additional drug therapy with rituximab or bortezomib was instituted depending upon initial response.

### Antibody Testing

The details of antibody testing performed at our center have been described previously^21^. Briefly, pre-transplant complement-dependent cytotoxicity (CDC) assays and three-color flow-cytometric cross matching (FCXM) were performed for all patients at the time of transplant. Donor-specific antibodies (DSA) were analyzed using the Luminex platform (Immucor Platform, San Diego, CA) with the use of an HLA phenotype panel (Lifematch Class I and Class II ID, Gen-Probe) and a single-antigen panel (Single Antigen Beads, Immucor Platform). Results of bead assays were measured as mean fluorescence intensity (MFI). For highly sensitized patients an MFI of >5000 was considered unacceptable while for de-novo kidney transplant patients an MFI of >10000 was considered unacceptable for kidney transplantation. Calculated Panel Reactive Antibody (cPRA) was calculated using CPRA calcoletor^22^.

### Lipidomic Analysis

#### Lipids extraction

Serial plasma samples were stored at -80°C prior to research use. Upon initiation of experiments, plasma samples were prepared for analysis using a HILIC-based UPLC ESI-MS/MS method. 50 µL of plasma was added to 750 µL of MTBE (methyl-tertiary butyl ether), containing 20 µL of SPLASH internal standards (SPLASH LIPIDOMIX Mass Spec Standard – Avanti 330707), and 160 µL of water. After centrifugation for 2 minutes at 12,300 rpm, 350 µL of supernatant was transferred to auto sampler vials and dried under vacuum. Dried extracts were re-suspended using 110 µL of a methanol:toluene (10:1, v/v) mixture containing CUDA (12-[[(cyclohexylamino) carbonyl] amino]-dodecanoic acid)at a final concentration of 50 ng/ml.

#### Lipidomic data acquisition via mass spectrometry

Samples were analyzed on a QTRAP 6500+, with Shimadzu Nexera UPLC. Analytes were separated on a Waters BEH HILIC 1.7 μm 2.1x150 mm column (column temperature = 30°C). Mobile phase A: 10 mM ammonium acetate (pH 8) in 95% ACN (acetonitrile). Mobile phase B: 10 mM ammonium acetate (pH 8) in 50% ACN. Gradient (B%) ramps from 0.1 to 20 in 10 mins; rises to 98 at 11 min, keeps for 2 mins, then drops back to 0.1 and maintains for 3 mins.

### Statistical Analysis

A comparison t-test analysis (FDR=0.05) was used to select group differences on the day of transplant. Lipids classes mean values were obtained by sum and average of all the lipids by each class. Linear Discriminant Analysis with regularized correction (RLDA) models for lipids and clinical parameters were created with stepwise forward method (Figure 1). Regression performance was estimated with R^2^, misclassification error and area under the ROC Curve (AUC). Estimates were validated with bootstrap coefficient interval (Figure 1). Predictors combined model was cross validated with Random Forest method, and the misclassification out-of-bag error (OOB error) was estimated and compared to the RLDA error for validation (Figure 1). Changes over time were also estimated with sparse partial least square method and group’s separation was validated with permutation test. T-test comparison of two time points for the same group and for comparing groups at matched time points. Data was analyzed with JMP Pro 13 and MetaboAnalyst 3.0. The statistical workflow is depicted in Figure 1.

**Figure 1:**
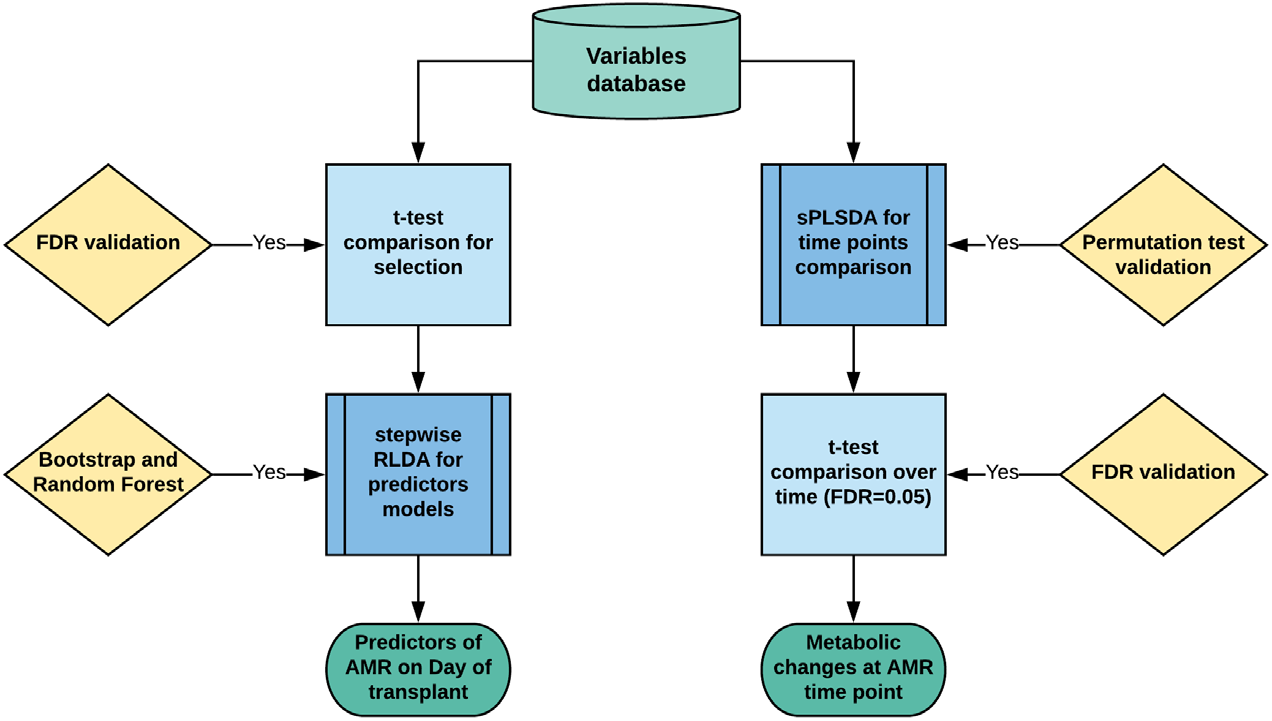
Statistical analysis workflow for the study. After data filtering and normalization a statistical workflow based on Regularized Linear Discriminant Analysis (RLDA) and Sparse Partial Least Square Discriminant Analysis (sPLSDA) was applied. Variables were selected by t-test with FDR=0.05 validation. RLDA on the day of transplant was used to identify predictor’s models of rejection on day of transplant with lipids, clinical parameters, and a combined model through stepwise forward method. Bootstrap and Random Forest were used as internal validation. sPLSDA at three different time points was used to identify and compare metabolic changes indicative of AMR. Permutation test was used as validation.

## RESULTS

### Clinical characteristics of the study population

Demographic comparison of the two groups in the day of transplant is described in Table 1. It revealed that there are significant differences for the following parameters for the non-rejection (SC) compared to the rejection (AMR) group. Patients in the AMR group were more likely to be female, re-transplants and had a higher degree of sensitization (higher cPRA) and presence of donor specific antibody (higher DSA) at the time of transplant. They were also more likely to have hyperlipidemia. There were no differences noted for age, race, weight, years on dialysis, type of dialysis, delayed graft function, or the presence or absence of diabetes mellitus.

**Table 1.** – Demographic Characteristics of the Patient Cohort. Categorical variables were analyzed with the Fisher’s exact test; Continuous data is presented as a mean of the group ± standard deviation and is analyzed by t-test. SD: Standard deviation; cPRA: calculated panel reactive antibody; DSA: donor specific antibody; GRF: glomerular filtration rate.

| Characteristic | SC | AMR | p-value |
| --- | --- | --- | --- |
| **N** | 29 (100%) | 16 (100%) |  |
| **Female Gender** | 4 (14%) | 11 (69%) | 0.005* |
| **Age, years (Mean±SD)** | 47±11 | 50±9 | 0.45 |
| **African-American Race** | 17 (59%) | 13 (81%) | 0.19 |
| **Pre-transplant Diabetes** | 10 (34%) | 8 (50%) | 0.35 |
| **Pre-transplant hyperlipidemia** | 7 (29%) | 16 (100%) | 0.04* |
| **Weight at Transplant, kg (Mean±SD)** | 85±21 | 82±14 | 0.6 |
| **Years on dialysis (Mean±SD)** | 2.9±1.9 | 4.3±4.1 | 0.26 |
| **Mode of dialysis** |  |  |  |
| **Hemodialysis** | 19 (65%) | 13 (81%) | 0.49 |
| **Peritoneal Dialysis** | 4 (14%) | 2 (12%) |  |
| **Preemptive transplant** | 6 (21%) | 1 (7%) |  |
| **Re-transplant** | 4 (14%) | 9 (56%) | 0.001* |
| **cPRA, % (Mean±SD)** | 9.8 (±29.4) | 40.8(±45.8) | 0.023* |
| **DSA** | 1 (3%) | 8 (50%) | <0.001* |
| **Kidney Donor Profile Index, %** | 52±27 | 54 ±32 | 0.89 |
| **Delayed Graft Function** | 13 (45%) | 7 (44%) | 1.00 |
| **GFR at 6 months post-transplant\*** | 67±22 | 61±23 | 0.37 |
| **GFR at 12 months post-transplant\*** | 68±19 | 58±22 | 0.11 |

### Circulating phospholipid concentrations was significantly different between SC and AMR prior to transplantation

A comparison of lipids classes on the day of transplant revealed PLs relative concentration differences between SC and AMR (Figure 2). Concentration of phosphatidylcholine (PC) was significantly diminished in AMR, while there was a trend for increased concentration of lysophosphatidylcholine (LPC). AMR group also showed significantly lower concentration of phosphatidylethanolamine (PE), lysophosphatidylethanolamine (LPE), plasmanylethanolamine (PE-O), and plasmenylethanolamine (PE-P). Although not statistically significant, there was also lower concentration of Phosphoglycerol (PG), lysophosphatidylglycerol (LPG), and sphingomyelin (SM). The activity of phospholipase A_2_ (PLA_2_) as a signal of increased metabolism was accessed by the ratio of phospholipids (PLs) to lysophospholipids (LPLs). AMR group showed decreased ratios of PC/LPC and PE/LPE indicating higher activity of PLA_2_ in this group on the day of transplant. This enzyme activity towards PLs degradation was evident for PE, showing more activity in the AMR.

**Figure 2:**
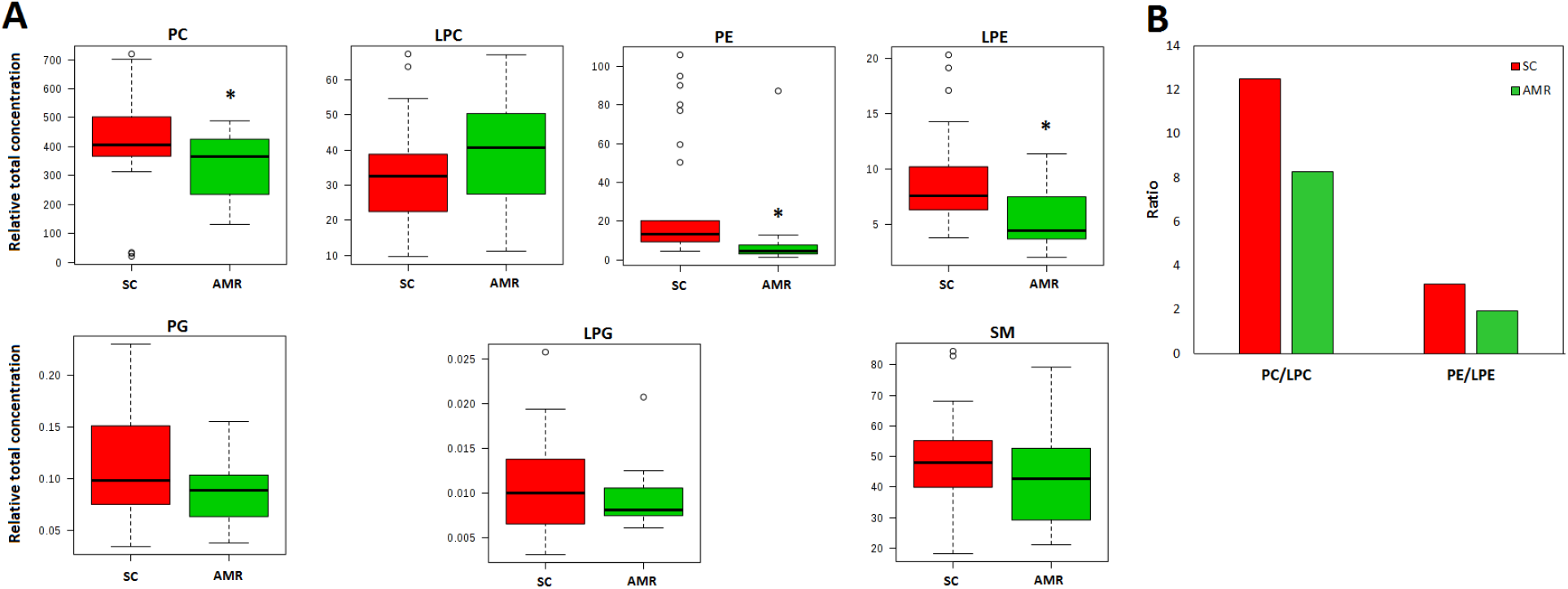
Significant differences are observed among phospholipids at T1 between SC and AMR. A) AMR group showed a significant lower concentration of PC, PE, and LPE. There was a trend towards higher levels of LPC in AMR. B) The ratio of PLs degradation to produce LPLs is an indication of PLA_2_ activity with lower values suggesting higher activity. AMR group presented lower ratio for both more PC/LPC and PE/LPE. Suspected outliers are indicated by open circles in the box plot. Green rectangles represent AMR and the Red rectangles represent SC. * indicates significant differences with p<0.05.

### Combined lipid and clinical parameters allow the prediction of rejection on the day of transplant

Our data so far demonstrated that there are significant differences in the lipidome between SC and AMR on the day of transplant. This led to the hypothesis that the lipidome or some combination of the lipidome and clinical parameters may enable the prediction of kidney rejection at time of transplant, allowing for risk stratification of graft recipients. To investigate this possibility, a stepwise regularized linear regression with only lipids, only clinical data and a merged clinical and lipid model was tested to see if the lipids alone or a combination of lipid and clinical variables would provide a model with high prediction accuracy (Table 2). The analysis identified 7 distinct lipids that discriminated between AMR and SC with 8.9% of the events misclassified, area under receiver operating characteristic curve (AUC) =0.95 (95%CI=0.84-0.98), R^2^=0.63 (95%CI=0.4-0.8). A clinical model using cPRA and DSA was inferior with 15.6% of the events misclassified, AUC=0.80 (95%CI=0.66-0.90), R^2^=0.36 (95%CI=0.16-0.57). Still using a stepwise selection approach, a combined model determined with 4 lipids plus DSA further reduced the misclassification events to 2.2% (Figure 3), and the AUC improved to 0.97 (95% CI=0.88-1.0), R^2^=0.81 (95%CI=0.49-0.96).

**Figure 3:**
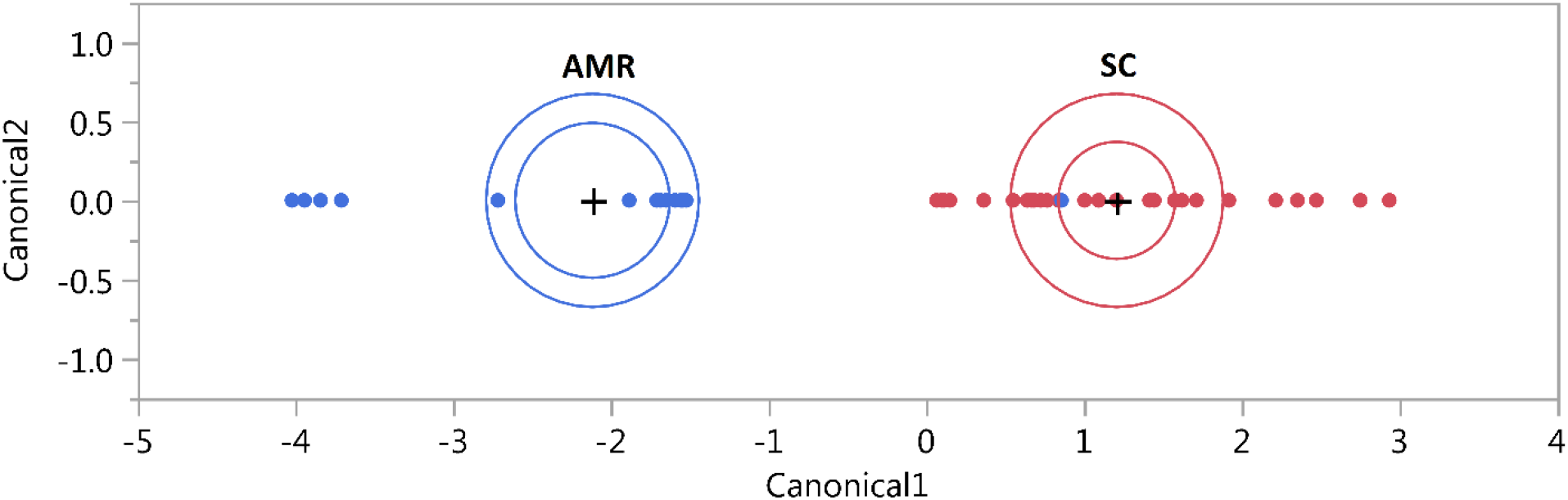
The RLDA model generated using 4 lipids and DSA demonstrate good separation between AMR and SC groups. The RLDA plot shows the clear separation of the patients in the two groups based on the Mahalanobis distance. This method determines whether the selected predictors can separate the distinct categories and reveals the presence of outliers in in the AMR and SC groups. Blue dot among the red dots indicates the one misclassified patient based in the predictive model. Internal ellipse indicates the 95% confidence region to contain the true mean of the group. External ellipse indicates the region estimated to contain 50% of group’ population.

**Table 2.** – Predictors of Rejection at the Time of Transplant. Bootstrap validation with 95% Confidence intervals is included for RLDA estimates and area under the curve (AUC). cPRA: Calculated Panel Reactive Antibody; DSA: donor specific antibodies; GFR: Estimated glomerular filtration rate (mL/min/1.73m2); SC: Stable Controls; AMR: Antibody-mediated Rejection; *statistically significant.

| Model | Predictors | R<sup>2</sup> | Misclassification<br>n | AUC |
| --- | --- | --- | --- | --- |
| **Only lipids** | PC (16:0/22:6) | 0.63 | 8.9% | 0.95 |
|  | PC (18:0/20:4) | (0.40 – 0.80) | (3.3 – 18.6) | (0.84 – 0.98) |
|  | PC (18:1/20:4) |  |  |  |
|  | LPE (16:0) |  |  |  |
|  | LPE (16:1) |  |  |  |
|  | LPE (20:4) |  |  |  |
|  | LPE (22:6) |  |  |  |
| **Only clinical** | cPRA | 0.36 | 15.9% | 0.80 |
|  | DSA | (0.16 – 0.57) | (7.4 – 29.2) | (0.66 -0.90) |
| **Merged models** | PC (18:0/20:4) | 0.81 | 2.3% | 0.97 |
|  | LPE (16:0) | (0.49 – 0.96) | (0.1 – 12.1) | (0.88 – 1.00) |
|  | LPE (20:4) |  |  |  |
|  | LPE (22:6) |  |  |  |
|  | DSA |  |  |  |

Further comparison of the 4 lipids predictors of kidney rejection between SC and AMR showed that these lipids are significantly decreased in AMR group, although in PC (18:0/20:4) plot it is possible to notice presence of outliers in both groups (Figure 4A). Random Forest method was used for statistical validation with 500 bootstrap samples, and the mean decrease accuracy test was used estimate the importance of each predictor to the validation model (Figure 4B). The result revealed that DSA is the more important biomarker of AMR at the day of transplant, and together with LPE (16:0) and PC (18:0/20:4) can discriminate AMR with very low error (2.2%). The statistical validation also revealed that exclusion of LPE (22:6) and LPE (20:4) of the model would not affect much the misclassification error, although in the RLDA modeling training, using the study population, the addition of these two lipids takes the model estimation from R^2^=0.75 to R^2^=0.81.

**Figure 4:**
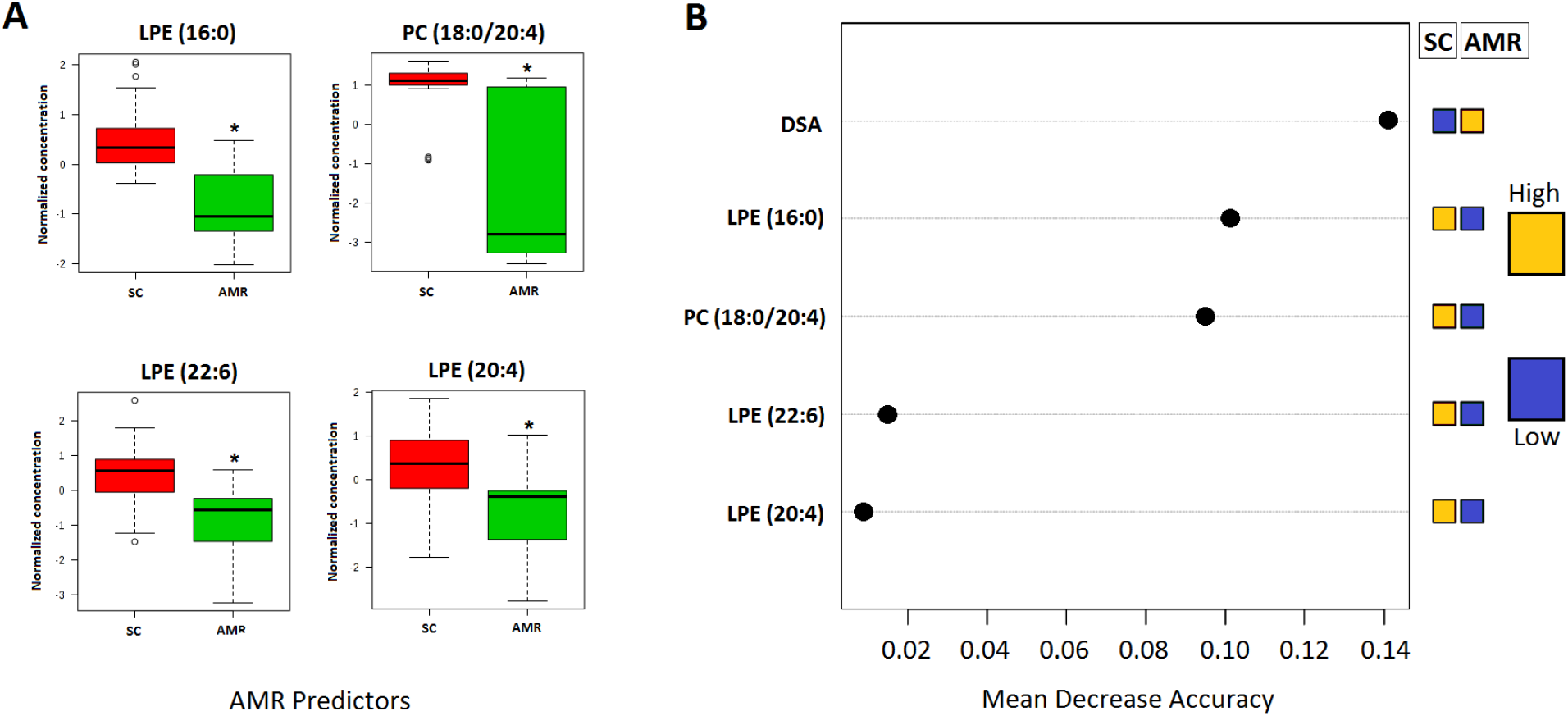
Predictors of AMR on the day of transplant and Random Forest statistical validation. A) Box plot of normalized concentration shows that AMR group has lower concentration of the lipids predictors. Suspected outliers are represented as open circles that appear outside the whiskers. The validation method showed that the predict model can discriminate SC and AMR in the day of transplant with 0.022 OOB error. The mean Decrease Accuracy method shows that DSA is the more important predictor, followed by LPE (16:0) and PC (18:0/20:4) and they independently could be used as biomarkers. The analysis also reveals that when considering these predictors as biomarkers, the inclusion of LPE (20:4) and LPE (22:6) does not add any predictive power, and rather must be use to compose the RLDA model. * indicates significant differences with p<0.01.

### Serial analysis of the lipidome over the course of one year identify time dependent lipid changes among patients with a favorable transplant outcome

Following the identification of the lipid differences at T0 and their ability to predict graft rejection in association with measured clinical parameters, we wished to investigate how the lipidome changes over time in patients with a favorable transplant outcome (SC). To achieve this end, lipid profiles were analyzed between serially collected samples at Day 0, 6 months and 12 months post-transplant (Figure 5). A sPLSDA analysis of the data revealed a statistically significant alteration in the metabolic profile at 6 months post-transplant compared to the day of transplant (Figure 5A). However, for the subsequent times from 6 months to 12 months, there was no significant change in the lipidomic profile (Figure 5B). The data was subjected to validation using the permutation test (Figure 5C) and showed significant metabolic difference (*p*= 0.034) from day of transplant to 6 months after transplantation.

**Figure 5:**
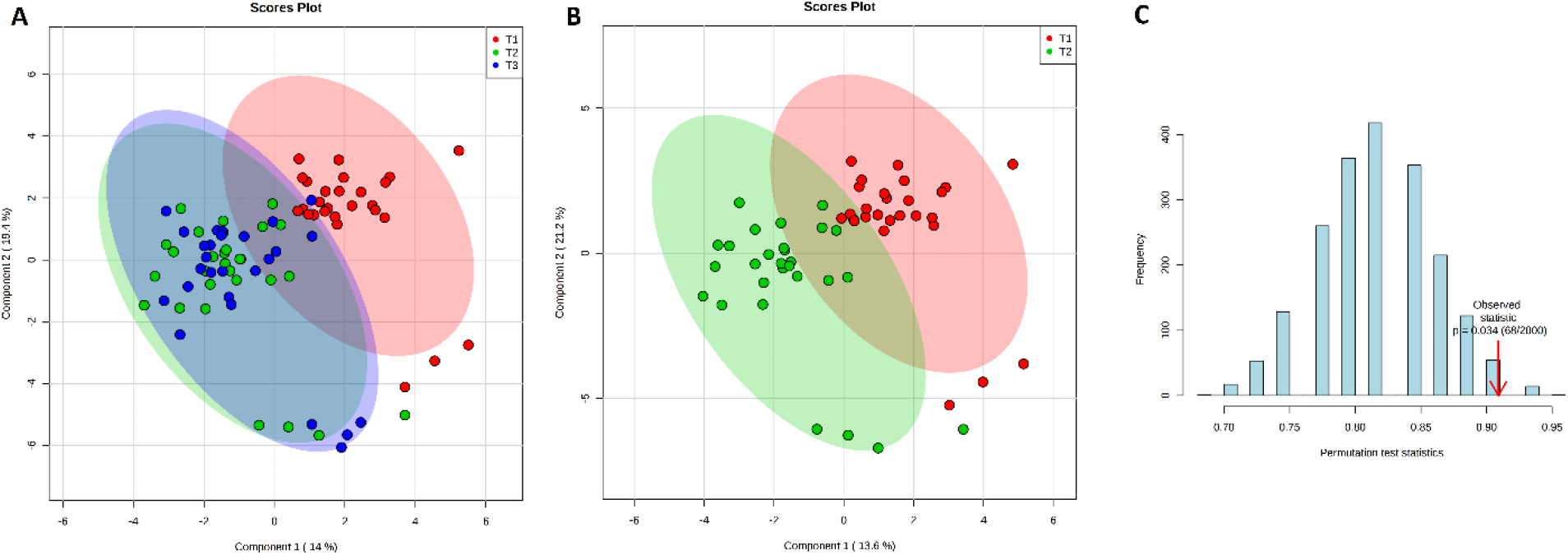
The lipidome of SC demonstrate clear differences between T1 and T2 but no differences between T2 and T3. A) The graphical distribution of T1 (shown in red), T2 (shown in green), and T3 (shown in blue) indicates that there is no difference between 6 months and 1-year post-transplant, after a metabolic shift from the day of transplant. B) The lipid difference is highlighted by the change in the first 6 months C) Permutation test was performed as a validation test to evaluate the statistical significance of the PLS-DA model separation from T1 to T2 (p=0.034). Ellipses represent the 95%CI of each time point.

Further investigation of the lipid differences between T1 and T2 identified 19 lipids that were significantly elevated at T2 compared to T1 among the SC. statistically relevant lipid changes that characterized the time dependent alteration to the lipidome among the SC (Figure 6). A majority of these lipids changes are LPC, with a few PC, one PE-O, two PE-P, and one PG.

**Figure 6:**
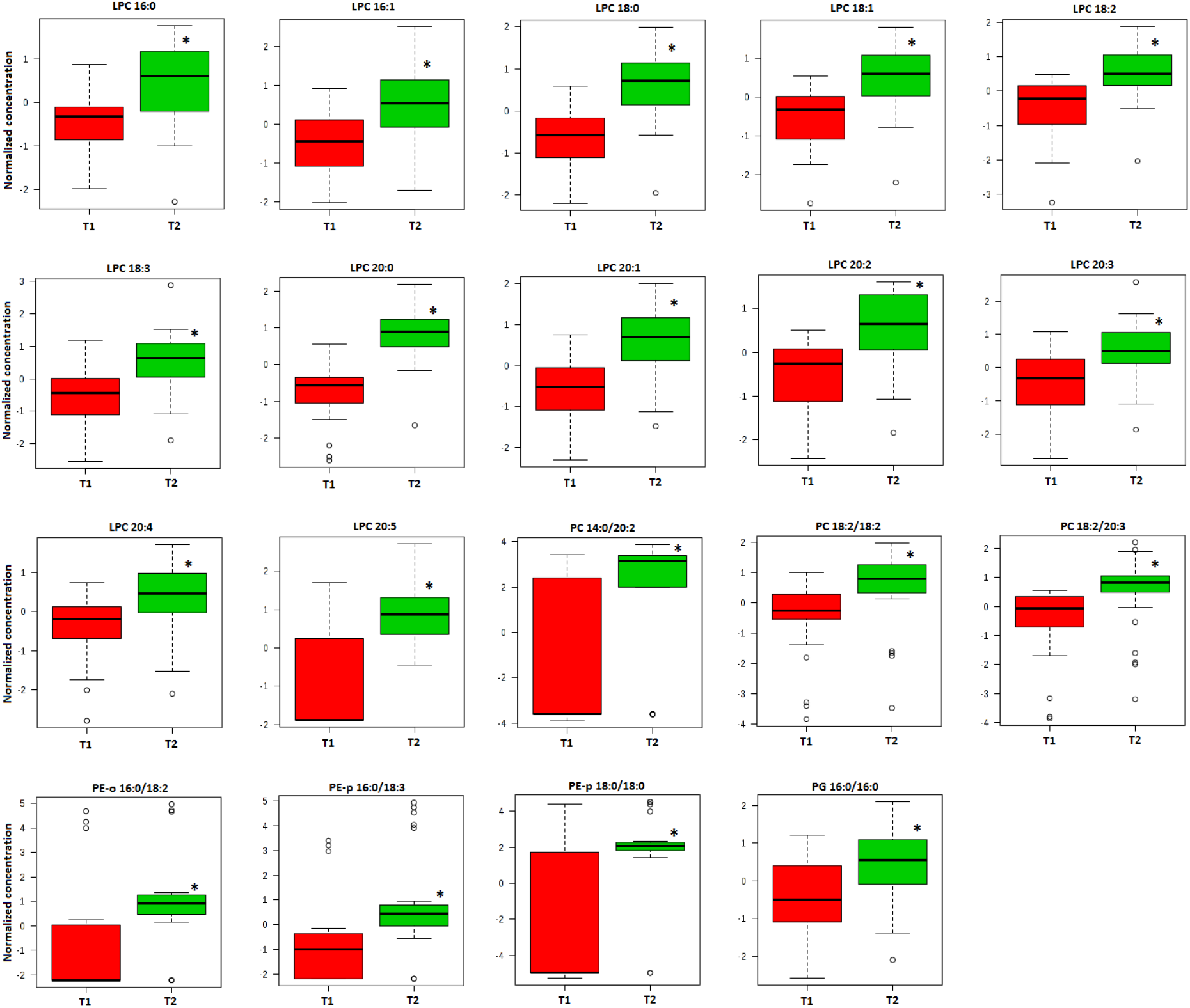
Specific lipids characterize the difference between T1 and T2 among SC patients. The levels of 19 different lipids are significantly elevated 6 months after transplantation. Most of the lipids are from LPC class and they contain saturated and saturated fatty acids. PCs, PE-O, PE-P and PG are also represented elevated after 6 months. * indicates significant differences with p<0.01.

### Serial analysis of the lipidome over the course of one year demonstrate no significant differences among the measured lipids among graft recipients with non-favorable outcomes

Following the identification of the longitudinal lipid trajectory among patients with favorable transplant outcomes, we investigated the trajectory of the lipidome pre-transplant to post transplant one year among the patients with non-favorable outcomes (AMR) (Figure 7). sPLSDA analysis of the data reveal that there was no significant alteration in the lipid profile at pre-rejection and post-rejection compared to the day of transplant (Figure 7a). While a slight change was observed from day of transplant to post-rejection (Figure 7B), validation analysis using permutation testing demonstrated this difference to be non-significant (p=0.869) (Figure 7C).

**Figure 7:**
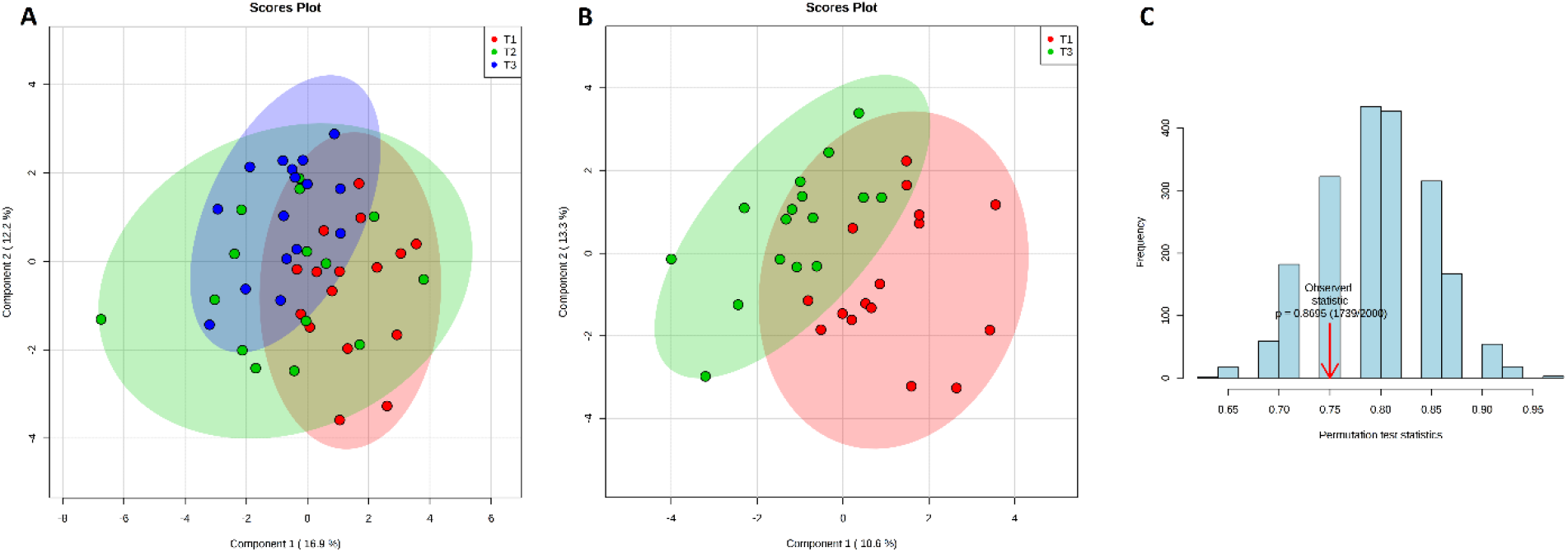
Contrary to SC patients, no statistically significant difference was observed in the T1 and T2 lipidome of AMR patients. A) The graphical distribution of T1 (shown in red), T2 (shown in green), and T3 (shown in blue) indicates that there is no difference over time, although a slight metabolic shift could be detected from the day of transplant to post-rejection. B) The plot of the slight metabolic difference from day of transplant to post-rejection highlight the overlap of the 95%CI of the two time points. C) Permutation test was performed as a validation test to evaluate if there is a statistical significance of the PLS-DA model separation from T1 to T2 (p=0.869). Ellipses represent the 95%CI of each time point.

### Significant lipid changes were also observed between patients favorable and non-favorable transplant outcomes at post-transplant, pre-rejection time point (T2)

As our data revealed that there were significant T1 vs T2 lipid differences between SC, but not in AMR, we further investigated the data to identify the exact differences in the lipidome between SC and AMR at T2. Any differences identified would indicate an alteration in the lipid metabolic environment at the time of rejection that would distinguish AMR from SC. Since there were no significant differences between T2 and T3 for SC group we chose to use SC at T2 (6 months post-transplant) to compare with AMT at T3 (time of AMR). The analysis revealed a panel of 13 lipids that were found to differentiate the two groups at T2 (Figure 8). As before, these 13 lipids were again comprised of LPE and PC species containing monounsaturated and polyunsaturated fatty acids, except for LPE (16:0). This data further confirms the presence of a sustained lipid metabolic difference between SC and AMR over time that distinguish between the patients with favorable and non-favorable transplant outcomes.

**Figure 8:**
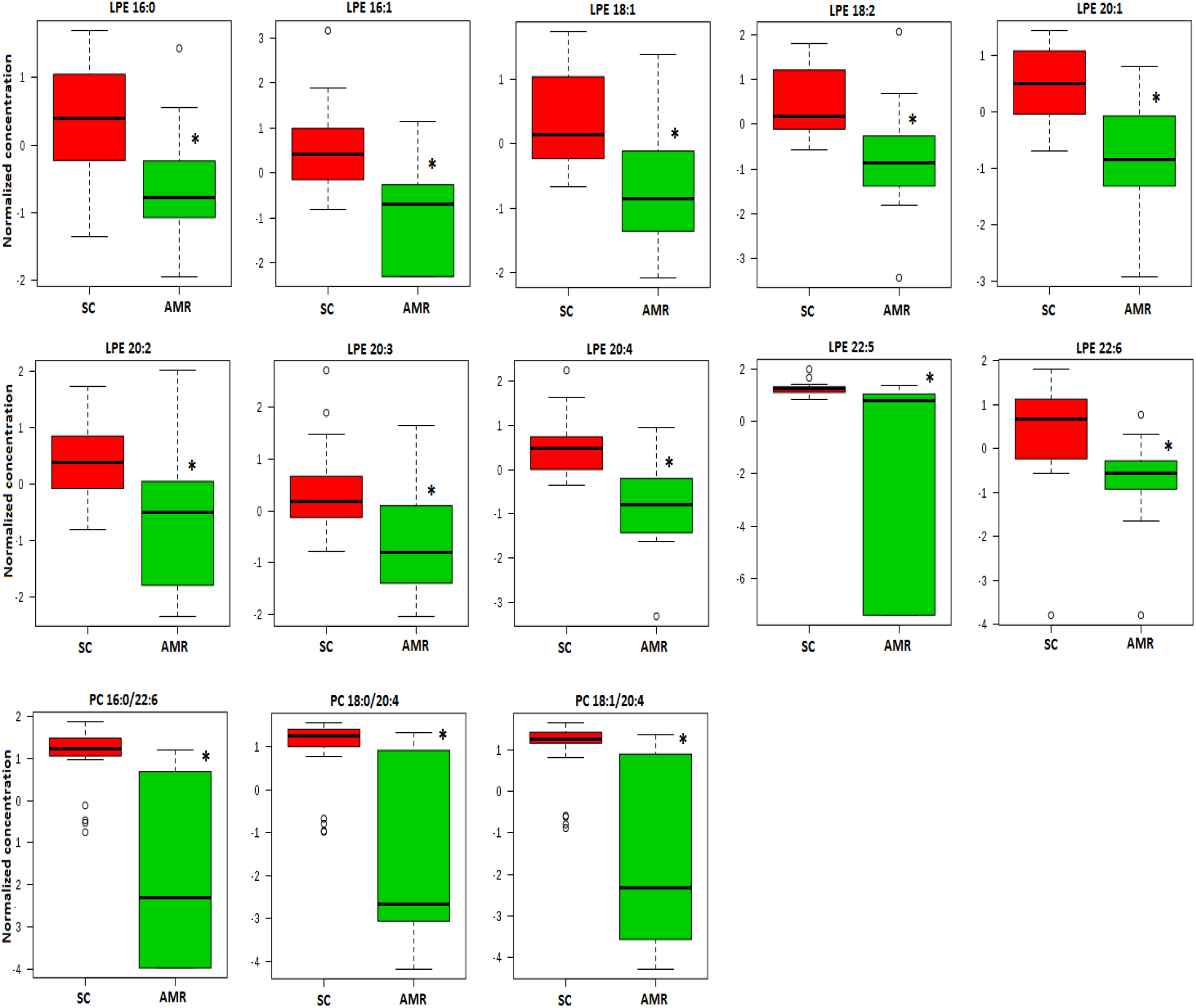
Specific lipids demonstrate significant differences between SC and AMR at T2. The metabolic changes observed in the day of transplant was sustained 6 months after transplant with lower LPE and PC species in AMR group. Except for LPE (16:0) all lipids contained monounsaturated and polyunsaturated fatty acids. SC group shown in red. AMR group shown in green. * indicates significant differences with p<0.01.

## DISCUSSION

This study highlights the lipidomic signature differences by comparing the lipid profiles of SC and AMR groups at various periods before and after kidney transplantation. To the best of our knowledge, this is the first study to evaluate the lipidomic profile longitudinally and predict the risk of developing kidney rejection. As we face a disorder with unique complexity and unclear pathogenesis, our study, among other previous studies^23^, has demonstrated that circulating lipid abnormalities have a role in the etiology of renal allograft rejection. The specific pattern of lipidomic abnormalities pre and post kidney transplantation, as well as their links to inflammation and rejection, are unclear. However, we explored the changes and possibilities of AMR compared SC group to better understand the processes and function of lipidomic in these individuals.

The comparison of SC and AMR groups on the day of transplant revealed that female gender, re-transplant, cPRA, DSA, and hyperlipidemia were statistically different. Moreover, we found DSA as the strongest predictor of AMR. These findings are consistent with Dunn *et al* where it was reported that DSA and female gender were risk factors for AMR, while cPRA was associated with cell-mediated rejection. However, re-transplant did not reach statistical significance^24^. Our finding of hyperlipidemia in the AMR group could be linked to the fact that hyperlipidemia is the most common form of dyslipidemia, a common complication in CKD patients, associated with the decline in kidney function, hypertriglyceridemia, low HDL, and low or normal LDL^25^.

The lipidomic profile that we observed between SC and AMR prior to transplantation revealed some circulating PLs s that are significantly different, which is mostly in agreement with other previously published studies suggesting PLs being the most valuable class of lipids altered in kidney transplant patients^26–29^. Modulation of PLs in CKD is well described in the literature. To illustrate, Anna Michalcyke *et al* showed that LPA concentrations were statistically significant higher in patients with kidney transplants than healthy individuals (without CKD)^27^. Additionally, the same study confirmed that patients with unfavorable renal outcomes, including patients with renal transplantation, are associated with lower LPC levels compared to healthy individuals^27^. Furthermore, an observational study of CKD patients with either glomerulonephritis or subacute/chronic tubulointerstitial injury compared to in healthy volunteers showed that changes in urinary phospholipids, especially LPCs, were associated as a result from proteinuria damaged kidney function or proteinuria induced hypoalbuminemia or lipotoxicity^26^. Additionally, this has been confirmed by Lewen Jia *et al* where plasma samples of patients with chronic glomerulonephritis and patients with CKD without renal replacement therapy were compared to healthy persons. This study’s showed that primary chronic glomerulonephritis and CRF had phospholipid metabolic abnormality, suggesting that nineteen phospholipid species were identified as possible biomarkers in plasma samples of chronic glomerulonephritis and CKD^28^. As our work demonstrates that there are alterations in PIs such as PEs, PCs, and LPEs, PIs might be potential biomarkers for kidney injury and prediction of allograft rejection. This showed the potential role of PLs in the pathogenesis and immune/inflammation in kidney disease. The presence of alterations PIs concentrations observed in patients may have clinical significance.

LPC has been associated with pro-inflammatory effects^30^, but there is not much information on the effects of LPE. Some studies suggest that LPEs could have possible protective effect on inflammation. Schober *et al* demonstrated that LPE generation from PE oxidation is primarily due to PLA_2_^31^. The effect of PLA_2_ is well known by presenting pro-inflammatory action in the hydrolysis of PC to produce LPC with atherosclerotic properties, as well as anti-inflammatory action in the hydrolysis of platelet-activating factor (PAF) and oxidized PLs^32^. The activity of PLA_2_ in our study was accessed by the ratio of PLs to LPLs, with the AMR group showing higher PLA_2_ activity, especially for degradation of PE to produce LPE. The use of PC/LPC ratio is used as an indication of inflammation and could indirectly represent the increased activity of PLA2 in inflammatory diseases^33, 34^.

We have demonstrated that associating lipidomic and clinical data for multivariate analysis has the potential to reveal metabolic features of CKD patients undergoing dialysis and higher inflammation associated with hemodialysis^35^. In our study, merging DSA and lipids revealed higher DSA and reduction of each of the four identified lipid biomarkers: PC (18:0/20:4), LPE (16:0), LPE (22:6), and LPE (20:4), one PC and three LPE species, in the AMR group could discriminate AMR with minimal error. Additionally, those four lipids predictors of kidney rejection between SC and AMR on the day of transplant were found to be also significantly different at post-transplant (T2) PC (18:0/20:4), LPE (16:0), LPE (22:6), and LPE (20:4). Statistical validation suggested that LPEs and PCs are potential biomarkers that could be further investigated in a clinical study.

We found infection and rejection risk indicators of lipids that allowed us to stratify patients based on their relative risk of renal transplantation outcomes. Our method might allow us to predict patients’ specific risk of developing allograft rejection before transplantation. This approach might be used in more clinical settings to enhance the survival and quality of life of the thousands of patients who receive renal transplants. This is considered the first step toward customized medical therapy following transplantation, which allows us to create a risk profile for allograft rejection and the consequences of under and overimmunosuppression. This strategy’s potential utility is widespread, ranging from the prediction of patients who might undergo kidney rejection, identification of patients who could benefit from immunosuppressive reduction or tolerance induction methods to the optimization of organ allocation systems^18^.

The lack of significant differences in the lipidomic profile of the subsequent times from 6 months to 12 months with a favorable transplant outcome, indicating a stabilization of the lipid changes that happen after transplant with the achievement of improved kidney function and possibly a reduced milieu of inflammation. On the other hand, there was no significant alteration in the lipid profile on pre-rejection and post-rejection compared to the day of transplant among patients with AMR. These findings indicate that as opposed to patients with favorable transplant outcomes (SC), in the case of patients with non-favorable transplant outcomes (AMR), the lipid profile observed pre-transplant was sustained over time. Additionally, various mechanisms involving chronic inflammation, humoral, and cellular immune reactions play essential roles in the immunopathogenesis of kidney rejections. In fact, Transplant rejection can be classified as hyper-acute, acute, or chronic. Hyper-acute rejection is usually caused by specific antibodies against the graft and occurs within minutes or hours after grafting. Acute rejection occurs days or weeks after transplantation and can be caused by specific lymphocytes in the recipient that recognize human leukocyte antigens in the tissue or organ grafted. Lastly, chronic rejection usually occurs months or years after organ or tissue transplantation^36^. Therefore, showing significant lipid changes could fail because of the heterogeneity of types of kidney rejection, and different patient subgroups encounter various risks of rejection and infection^34^. Each type of allograft rejection has different mechanisms, so each type would likely have a different lipidomic profile and thus be difficult to evaluate unless large sample sizes of each type of kidney rejection are obtained^37^.

Identification of the factors that cause inflammation/immune response following solid organ implantation might lead to a better understanding of the underlying pathophysiology and new therapies for reducing inflammation inside the transplant, allowing the transplant to be treated before being implanted into the recipient. Inflammation resolution in renal transplantation is somewhat unknown territory. It may be possible to reduce the detrimental inflammation that occurs during transplantation by identifying the mechanisms that promote inflammation resolution. Approaches to successfully reduce local inflammation inside renal transplants, using lipidomic in combination with efforts to advance inflammation resolution, may improve the efficacy of renal transplantation, a vital treatment for end renal failure^38^.

Our results suggest that a lack of anti-inflammatory protection in patients on the day of transplant is a risk of rejection. No relevant changes occurred for AMR until rejection, confirming that the metabolic profile predicting AMR persisted after transplantation. Accordingly, over time comparison of SC and AMR showed that the difference in LPE level and PC was kept after 6 months representing the metabolic difference between rejection and non-rejection. The presence of monounsaturated and polyunsaturated fatty acids in PLs is also an indication that their low plasma content is a risk factor for kidney health^39^. In contrast, the elevation of LPC, PC, PE-O, PE-P, and PG after 6 months in the SC group implies that restauration of PLs content is a result of successful transplantation. Indeed, some studies have shown that elevation of polyunsaturated fatty acids presents a lower risk of developing end-stage renal disease^40^, as well as higher survival rates after kidney transplantation^41^. In a human study comparing health controls and CKD patients, Reis *et al* found that the content of total PC and Ceramides were decreased as well as the ratio of LPC/LPE^41^. In a study comparing CKD progressors patients compared to control patients, Afshinnia *et al* reported that CKD progressors had lower Cholesteryl ester (CE), diacylglycerols (DG), PC, plasmenylcholine (PC-P), PE-P, and phosphatidic acid (PA) while presenting elevated PE and monoacylglycerols (MAG)^42^. This suggests that CKD progressors with a decrease of longer acyl chains and polyunsaturated lipids could benefit from the effects of polyunsaturated fatty acid supplementation, as some studies reveal. In our study, although both groups represent progressive CKD patients, the SC group had higher PC and LPE than AMR and a trend for lower LPC.

The intrarenal renin-angiotensin system (RAS) has been postulated to have a role in the onset and progression of allograft damage, in addition to immunologic and inflammation pathways. The involvement of the intrarenal RAS in the pathogenesis of hypertension and renal damage has piqued researchers’ interest in recent years^43^. Hypertension (HT) is a common complication in kidney transplant patients, and it is a serious complication that can have a negative impact on patient and graft survival. Chronic allograft damage and graft failure have been observed as a result of HT. Despite the knowledge that HT is a manageable risk factor, kidney transplant recipients experience poor blood pressure management. Thus, HT is a significant risk factor for both graft and patient survival following transplantation^44^. HT, which is characterized by multiple changes in the structure and function of the cell membrane, such as changes in membrane microviscosity, receptor function, signal transduction, ion transport, calcium mobilization, intracellular pH regulation, etc, is frequently linked to serious metabolic abnormalities, including lipid metabolism. For example, alterations in membrane lipid composition due to extensive interchange between circulating and membrane lipids, as well as aberrant cellular lipid production and metabolism, are reflected in the changed membrane microviscosity. Lipids, as a component of the cell membrane, play a critical role in the control of the aforementioned membrane characteristics^45–47^. In fact, Phospholipids such as phosphatidylcholines (PC), phosphatidylethanolamines (PE), lysophosphatidylcholines (LPC), PE-based plasmalogens (PLPE), ceramides (CERs), and sphingomyelin (SPM) all play a role in the bilayer of (blood) cell membranes^47^. Understating the relationship between the lipid circulation and the renin-angiotensin system may reveal potential molecular targets that are involved in allograft failure as a result of HT. Phospholipids, which can give insight into the pathophysiology of hypertension, are measured by Jun Liu *et al*. In this study, eight circulating phospholipids have been discovered and linked to blood pressure that can predict the onset of hypertension. Six phosphatidylethanolamines (PE 38:3, PE 38:4, PE 38:6, PE 40:4, PE 40:5, and PE 40:6), as well as two phosphatidylcholines (PC 32:1 and PC 40:5), were shown to predict the occurrence of hypertension^48^. These findings are aligned with our specific significant lipids (figures 6 and 8). This suggests that phospholipid metabolites in the bloodstream might provide insight into blood pressure control, as well as a variety of viable hypotheses for future research indicating the role of the onset and progression of allograft rejection.

The study’s main limitation was that our pathological subgroups of kidney rejection were limited due to the small number of patients in the cohort. Second, Cellular rejection and systemic inflammatory illnesses, for example, were excluded due to a lack of individuals with those symptoms. The addition of these pathologic symptoms, which are associated with elevated inflammation, may have an impact on our lipidomic risk stratification and prediction of kidney rejection. Cross-validation at multiple centers with higher patient numbers and disease subgroups is needed to confirm our preliminary research results.

## CONCLUSION

Our study, for the first time identifies the lipid differences pre-transplant and post-transplant, pre-rejection that distinguish kidney transplant patients with favorable transplant outcomes (SC) and a major form of non-favorable transplant outcomes (AMR). We further demonstrate that unlike SC patients that demonstrate a dynamic longitudinal lipid change, AMR patients maintain a relatively unchanging lipid profile over time with respect to the measured lipids. In addition, we demonstrate for the first time the potential for risk stratification of kidney transplant patients on the day of transplant with respect to the potential for the onset of AMR. Lastly, we identify some potential target lipids that could involve in some inflammatory processes of kidney rejection. Following validation in a larger cohort, these findings have the potential to alter the current paradigm of post-transplant monitoring and treatment of these patients via an evidenced-based risk stratification strategy and thereby vastly improving the success of kidney transplantation.

## DISCLOSURE

There are no conflicts of interest to report for any of the authors.

## ACKNOWLEDGEMENT

We would like to acknowledge the contributions to this research by Drs. Jeffrey Stern, Naren Gajenthra Kumar, Pamela Kimball, Anne King, Dhiren Kumar and Marlon Levy. Research reported in this publication was supported by research grants from National Institutes of Health under grant numbers HD087198 (to DSW). The content is solely the responsibility of the authors and does not necessarily represent the official views of the National Institutes of Health. This work also received support via a Young Investigator Award from SCIEX for clinical lipidomic research (DSW).

## Notes

### Competing Interest Statement

The authors have declared no competing interest.

### Summary of Updates

The introduction has been modified to be in line with critiques obtained from reviewers.

